## Supplementary File for "*Fruit-In-Sight:* a deep learning-based framework for secondary metabolite class prediction using fruit and leaf images"

Supplementary File 1: Prediction combinations under the multi-analyte framework to be used for low (A) and high (B) fruit azadirachtin class prediction.

A

| $A_{fruit}$ | $D_{fruit}$ | $E_{fruit}$ | $N_{fruit}$ | $S_{fruit}$ | $A_{leaf}$ | $D_{leaf}$ | $E_{leaf}$ | $N_{leaf}$ | $S_{leaf}$ |
|---|---|---|---|---|---|---|---|---|---|
| l | x | x | x | h | x | l | x | x | l |
| l | x | x | x | x | l | l | h | l | x |
| l | x | x | x | x | l | l | l | x | l |
| l | x | x | x | x | l | l | x | l | l |
| l | x | x | x | x | x | l | l | h | l |
| l | l | x | x | x | l | l | x | l | x |
| l | x | l | x | x | l | l | x | l | x |
| l | x | l | x | x | l | l | x | x | l |
| l | x | l | x | x | l | x | h | l | x |
| l | x | l | x | x | x | l | x | h | l |
| l | x | l | l | x | l | l | x | x | x |
| l | x | l | l | x | l | x | h | x | x |
| l | x | l | h | x | l | x | x | l | x |
| l | x | l | x | l | l | l | x | x | x |
| l | x | l | x | l | l | x | h | x | x |
| l | x | h | x | l | x | x | l | h | x |
| l | x | x | l | x | l | l | x | l | x |
| l | x | x | l | x | x | l | l | x | l |
| l | x | x | x | l | l | l | x | l | x |
| x | l | l | h | x | l | x | x | l | x |
| x | x | l | x | x | l | l | l | h | x |
| l | l | x | x | x | l | l | l | h | x |
| l | l | x | x | x | x | l | h | l | l |
| l | l | l | x | x | l | l | l | x | x |
| l | l | l | x | x | l | x | x | l | l |
| l | l | l | x | x | x | l | h | x | l |
| l | l | h | x | x | x | h | x | h | h |
| l | l | l | x | x | x | x | h | l | l |
| l | l | l | l | x | l | x | x | x | l |
| l | l | l | l | x | x | l | x | x | l |
| l | l | h | h | x | x | x | x | h | h |
| l | l | l | l | x | x | x | x | l | l |
| l | l | l | x | l | x | x | h | l | x |
| l | l | x | l | x | l | l | l | x | x |
| l | l | x | l | x | l | x | l | x | l |
| l | l | x | h | x | x | h | x | h | h |
| l | l | x | l | x | x | l | x | l | l |
| l | l | x | l | x | x | x | l | l | l |
| l | l | x | l | h | l | l | x | x | x |
| l | l | x | l | l | x | x | x | l | l |
| l | l | x | x | l | l | l | l | x | x |
| l | l | x | x | h | x | h | x | h | h |
| l | x | l | h | x | l | l | l | x | x |
| l | x | l | h | h | l | x | l | x | x |
| l | x | l | l | l | l | x | x | x | l |
| l | x | l | x | l | l | x | x | l | l |
| l | x | h | x | l | x | h | h | x | l |
| l | x | x | h | l | l | l | l | x | x |

| Legend | | |
|---|---|---|
| l | : | low |
| h | : | high |
| x | : | low or high, doesnt matter |
| A | : | azadirachtin |
| D | : | deacetyl salannin |
| E | : | nimbolide |
| N | : | nimbin |
| S | : | salannin |

| | | | | | | | | | |
|---|---|---|---|---|---|---|---|---|---|
| l | x | x | l | l | l | l | l | x | x |
| l | x | x | l | l | l | x | l | x | l |
| l | x | x | l | l | l | x | x | l | l |
| l | x | x | l | l | x | x | l | l | l |
| l | x | x | x | l | l | l | l | h | x |
| x | l | l | x | x | x | l | l | h | l |
| x | l | x | l | x | l | l | l | h | x |
| x | x | x | l | l | l | l | l | h | x |
| x | l | x | x | l | l | l | l | h | l |
| l | x | x | x | x | l | l | x | l | x |
| l | x | l | x | x | l | l | x | x | x |
| l | x | x | x | x | l | l | l | h | x |
| l | l | x | x | x | l | l | x | x | h |
| l | l | x | x | x | l | l | x | x | l |
| l | l | x | x | x | l | x | l | l | x |
| l | l | x | x | x | x | l | l | h | x |
| l | l | x | x | x | x | x | h | l | l |
| l | l | h | x | x | l | l | x | x | x |
| l | l | h | x | x | l | x | l | x | x |
| l | h | l | x | x | l | x | x | x | h |
| l | l | h | x | x | x | h | x | h | x |
| l | l | h | x | x | x | x | l | h | x |
| l | l | l | x | x | x | x | h | l | x |
| l | l | h | x | x | x | x | l | x | l |
| l | l | l | x | x | x | x | h | x | l |
| l | l | h | x | x | x | x | x | h | h |
| l | l | h | h | x | x | l | x | x | x |
| l | l | h | h | x | x | x | l | x | x |
| l | l | h | h | x | x | x | x | h | x |
| l | l | h | h | x | x | x | x | x | h |
| l | l | h | h | l | x | x | x | x | x |
| l | l | l | h | h | x | x | x | x | x |
| l | h | l | x | l | l | x | x | x | x |
| l | l | l | x | h | l | x | x | x | x |
| l | l | l | x | h | x | x | x | x | l |
| l | l | x | h | x | l | l | x | x | x |
| l | l | x | l | x | l | l | x | x | x |
| l | l | x | h | x | h | x | x | h | x |
| l | l | x | h | x | l | x | x | l | x |
| l | l | x | h | x | l | x | x | x | h |
| l | l | x | h | x | x | l | x | h | x |
| l | l | x | h | x | x | l | x | x | h |
| l | l | x | l | x | x | x | l | x | l |
| l | l | x | h | x | x | x | x | h | h |
| l | l | x | l | x | x | x | x | l | l |
| l | l | x | h | h | x | l | x | x | x |
| l | h | x | x | l | l | h | x | x | x |
| l | l | x | x | h | l | l | x | x | x |
| l | l | x | x | l | l | l | x | x | x |
| l | l | x | x | h | x | h | x | h | x |
| l | x | l | x | x | l | x | h | x | l |
| l | x | l | x | x | l | x | x | l | l |
| l | x | l | x | x | x | l | l | h | x |

| | | | | | | | | | |
|---|---|---|---|---|---|---|---|---|---|
| l | x | h | l | x | l | x | l | x | x |
| l | x | l | l | x | l | x | x | x | l |
| l | x | l | l | x | x | l | x | x | l |
| l | x | h | h | x | x | x | l | h | x |
| l | x | h | l | x | x | x | l | x | l |
| l | x | h | x | l | l | x | l | x | x |
| l | x | l | x | h | l | x | l | x | x |
| l | x | h | x | l | x | h | x | h | x |
| l | x | h | x | l | x | h | x | x | l |
| l | x | x | h | x | l | l | x | x | l |
| l | x | x | l | x | l | l | x | x | l |
| l | x | x | l | x | l | x | l | x | l |
| l | x | x | h | x | x | l | l | h | x |
| l | x | x | l | x | x | l | x | l | l |
| l | x | x | l | x | x | x | l | l | l |
| l | x | x | h | l | l | l | x | x | x |
| l | x | x | l | h | l | l | x | x | x |
| l | x | x | l | l | l | l | x | x | x |
| l | x | x | h | h | l | x | l | x | x |
| l | x | x | h | l | l | x | x | l | x |
| l | x | x | h | l | l | x | x | x | h |
| l | x | x | l | l | l | x | x | x | l |
| l | x | x | h | l | x | x | x | h | h |
| l | x | x | x | l | l | l | x | x | h |
| l | x | x | x | l | l | x | l | l | x |
| l | x | x | x | l | l | x | x | l | l |
| l | x | x | x | l | x | l | l | h | x |
| x | l | l | x | x | x | l | l | h | x |
| x | l | x | h | x | l | h | x | x | h |
| x | x | l | x | x | l | l | x | h | l |
| x | x | l | x | x | x | l | l | h | l |
| x | x | l | h | x | l | l | x | l | x |
| x | x | l | l | x | l | l | x | h | x |
| x | x | l | h | h | l | x | l | x | x |
| x | x | l | x | h | l | l | x | h | x |
| x | x | l | x | l | l | l | x | h | x |
| x | x | l | x | h | l | x | l | h | x |
| x | x | x | l | x | l | l | l | h | x |
| l | l | x | x | x | x | h | h | h | h |
| l | l | h | x | x | x | l | x | l | l |
| l | l | l | x | l | x | l | h | x | x |
| l | h | h | x | l | x | x | h | x | l |
| l | h | l | x | h | x | x | x | h | h |
| l | l | x | x | l | l | x | h | x | l |
| l | l | x | x | l | x | l | h | l | x |
| l | l | x | x | h | x | h | h | x | h |
| l | l | x | x | h | x | x | h | h | h |
| l | x | l | l | l | x | l | l | x | x |
| l | x | l | h | l | x | l | x | h | x |
| l | x | h | h | l | x | x | x | h | l |
| l | x | h | l | l | x | x | x | l | l |
| x | l | x | x | x | l | l | l | h | l |
| x | l | l | h | x | l | l | l | x | x |

| | | | | | | | | | |
|---|---|---|---|---|---|---|---|---|---|
| x | l | h | h | x | x | h | x | h | h |
| x | l | l | l | l | x | x | l | x | l |
| x | l | x | h | x | x | h | h | h | h |
| x | l | x | x | l | l | l | l | h | x |
| x | l | x | x | h | l | h | x | h | h |
| x | x | l | h | x | l | l | l | x | l |
| x | x | l | h | l | l | l | l | x | x |
| x | x | l | l | l | x | l | l | x | l |
| x | x | x | x | l | l | l | l | h | l |
| l | l | l | l | l | x | l | x | l | x |
| x | l | l | x | l | l | l | l | x | l |
| x | l | l | x | l | l | x | l | l | l |
| l | x | x | x | x | l | l | x | x | l |
| l | x | x | x | x | x | l | l | h | x |
| l | x | x | x | x | x | l | x | h | l |
| l | l | x | x | x | l | l | x | x | x |
| l | l | h | x | x | x | x | l | x | x |
| l | l | h | h | x | x | x | x | x | x |
| l | h | x | x | l | l | x | x | x | x |
| l | l | x | x | h | x | x | x | x | l |
| l | x | l | x | x | l | x | h | x | x |
| l | x | h | x | x | x | l | x | x | l |
| l | x | h | x | x | x | x | l | h | x |
| l | x | h | x | x | x | x | l | x | l |
| l | x | h | x | l | l | x | x | x | x |
| l | x | l | x | h | l | x | x | x | x |
| l | x | h | x | l | x | l | x | x | x |
| l | x | h | x | l | x | x | l | x | x |
| l | x | h | x | l | x | x | x | x | l |
| l | x | l | x | h | x | x | x | x | l |
| l | x | x | l | x | l | l | x | x | x |
| l | x | x | h | x | l | x | x | l | x |
| l | x | x | l | x | x | l | x | x | l |
| l | x | x | x | l | l | l | x | x | x |
| l | x | x | x | h | l | x | l | x | x |
| l | x | x | x | l | l | x | h | x | x |
| l | x | x | x | l | l | x | x | l | x |
| l | x | x | x | l | l | x | x | x | h |
| x | l | x | h | x | l | x | x | x | h |
| x | x | l | x | x | l | l | x | h | x |
| x | x | l | x | x | x | l | l | h | x |
| x | x | l | h | x | l | x | x | l | x |
| x | x | x | h | l | l | x | x | x | h |
| x | x | x | x | x | l | l | l | h | l |
| l | x | x | x | x | l | l | x | h | h |
| l | x | x | x | x | l | x | h | l | l |
| l | x | x | x | x | l | x | l | h | l |
| l | x | x | x | x | l | x | l | l | l |
| l | x | x | x | x | x | l | h | l | l |
| l | l | x | x | x | l | x | h | l | x |
| l | l | x | x | x | l | x | x | l | h |
| l | l | x | x | x | l | x | x | l | l |
| l | l | x | x | x | x | l | h | l | x |

| | | | | | | | | | |
|---|---|---|---|---|---|---|---|---|---|
| l | l | x | x | x | x | l | h | x | l |
| l | l | x | x | x | x | l | l | x | l |
| l | l | x | x | x | x | l | x | l | l |
| l | l | x | x | x | x | x | l | l | l |
| l | h | l | x | x | l | x | x | h | x |
| l | h | l | x | x | l | x | x | l | x |
| l | l | h | x | x | l | x | x | h | x |
| l | l | l | x | x | l | x | x | l | x |
| l | l | h | x | x | l | x | x | x | h |
| l | l | l | x | x | l | x | x | x | l |
| l | l | h | x | x | x | h | x | x | h |
| l | l | l | x | x | x | l | x | x | l |
| l | l | l | x | x | x | x | x | l | l |
| l | h | l | h | x | l | x | x | x | x |
| l | l | l | l | x | x | l | x | x | x |
| l | l | l | h | x | x | x | h | x | x |
| l | l | l | l | x | x | x | x | x | l |
| l | h | h | x | l | x | h | x | x | x |
| l | l | h | x | l | x | h | x | x | x |
| l | l | l | x | h | x | h | x | x | x |
| l | h | h | x | l | x | x | x | h | x |
| l | h | l | x | h | x | x | x | h | x |
| l | l | h | x | h | x | x | x | h | x |
| l | l | l | x | h | x | x | x | h | x |
| l | l | x | l | x | l | x | l | x | x |
| l | l | x | l | x | l | x | x | x | l |
| l | l | x | h | x | x | x | h | h | x |
| l | l | x | h | x | x | x | l | h | x |
| l | l | x | h | x | x | x | h | x | h |
| l | l | x | h | h | l | x | x | x | x |
| l | l | x | l | h | l | x | x | x | x |
| l | l | x | h | l | x | x | h | x | x |
| l | l | x | h | h | x | x | x | h | x |
| l | l | x | h | l | x | x | x | h | x |
| l | l | x | h | h | x | x | x | x | h |
| l | l | x | x | h | l | x | x | h | x |
| l | l | x | x | h | x | h | x | x | h |
| l | l | x | x | h | x | x | h | h | x |
| l | l | x | x | l | x | x | h | l | x |
| l | l | x | x | l | x | x | h | x | l |
| l | l | x | x | h | x | x | x | h | h |
| l | l | x | x | l | x | x | x | l | l |
| l | x | l | x | x | l | x | l | l | x |
| l | x | l | x | x | l | x | x | h | l |
| l | x | l | x | x | l | x | x | l | h |
| l | x | l | x | x | x | l | h | x | l |
| l | x | l | x | x | x | l | l | x | l |
| l | x | l | x | x | x | x | h | l | l |
| l | x | l | h | x | l | x | l | x | x |
| l | x | l | l | x | l | x | x | h | x |
| l | x | l | h | x | l | x | x | x | h |
| l | x | l | h | x | l | x | x | x | l |
| l | x | l | h | x | x | l | l | x | x |

| | | | | | | | | | |
|---|---|---|---|---|---|---|---|---|---|
| l | x | l | h | x | x | l | x | h | x |
| l | x | l | l | x | x | l | x | h | x |
| l | x | l | h | l | l | x | x | x | x |
| l | x | l | h | h | x | x | l | x | x |
| l | x | l | h | l | x | x | h | x | x |
| l | x | h | h | l | x | x | x | h | x |
| l | x | h | x | l | h | h | x | x | x |
| l | x | l | x | l | l | x | x | x | l |
| l | x | l | x | l | x | l | x | h | x |
| l | x | l | x | l | x | l | x | x | h |
| l | x | l | x | l | x | x | h | l | x |
| l | x | x | h | x | l | x | l | h | x |
| l | x | x | l | x | l | x | l | l | x |
| l | x | x | l | x | l | x | x | h | l |
| l | x | x | l | x | l | x | x | l | h |
| l | x | x | l | l | l | x | l | x | x |
| l | x | x | l | l | x | l | l | x | x |
| l | x | x | l | l | x | l | x | l | x |
| l | x | x | h | l | x | l | x | x | h |
| l | x | x | l | l | x | x | l | x | l |
| l | x | x | l | l | x | x | x | l | l |
| l | x | x | x | h | l | l | x | h | x |
| l | x | x | x | l | x | x | h | l | l |
| x | l | x | x | x | l | l | l | h | x |
| x | l | x | x | x | l | l | x | h | h |
| x | l | h | x | x | l | x | x | h | h |
| x | l | l | x | x | x | l | x | h | l |
| x | l | l | h | x | l | x | l | x | x |
| x | l | l | l | x | x | l | x | h | x |
| x | l | l | h | h | l | x | x | x | x |
| x | l | l | x | l | x | l | x | h | x |
| x | l | x | l | x | l | l | x | h | x |
| x | l | x | l | x | l | x | l | h | x |
| x | l | x | h | x | x | h | x | h | h |
| x | l | x | h | h | l | h | x | x | x |
| x | l | x | x | h | l | h | x | x | h |
| x | x | l | x | x | l | x | h | l | l |
| x | x | l | x | x | l | x | l | h | l |
| x | x | l | h | x | l | l | x | x | l |
| x | x | l | h | x | l | x | l | h | x |
| x | x | l | h | l | l | l | x | x | x |
| x | x | l | l | l | x | l | x | h | x |
| x | x | x | h | x | l | l | l | x | l |
| x | x | x | h | l | l | l | x | l | x |
| x | x | x | l | l | l | l | x | h | x |
| x | x | x | l | l | l | x | l | h | x |
| x | x | x | x | l | l | l | l | h | x |
| x | x | x | x | l | l | l | x | l | l |
| l | h | x | x | x | l | h | x | h | h |
| l | h | h | x | x | x | h | h | x | l |
| l | l | l | h | x | x | h | x | h | x |
| l | h | l | h | x | x | x | x | h | h |
| l | h | l | h | h | x | x | x | x | h |

| | | | | | | | | | |
|---|---|---|---|---|---|---|---|---|---|
| l | h | l | x | h | x | h | x | x | h |
| l | l | x | x | l | l | x | x | h | l |
| l | x | l | h | x | x | x | h | h | h |
| l | x | l | l | l | x | l | h | x | x |
| l | x | l | h | h | x | x | x | h | h |
| l | x | h | x | h | l | l | h | x | x |
| l | x | l | x | h | x | h | h | x | h |
| l | x | l | x | h | x | h | x | h | h |
| l | x | l | x | h | x | x | h | h | h |
| l | x | l | x | l | x | x | l | l | l |
| l | x | x | h | x | h | h | h | h | x |
| l | x | x | h | x | l | h | x | h | h |
| l | x | x | h | l | x | h | h | h | x |
| l | x | x | h | l | x | x | h | h | l |
| l | x | x | x | h | l | h | x | h | h |
| x | l | l | x | x | l | l | l | x | l |
| x | l | l | x | x | l | x | l | l | l |
| x | l | l | l | x | l | x | l | x | l |
| x | l | l | h | h | x | h | x | h | x |
| x | l | h | h | h | x | x | x | h | h |
| x | l | l | x | l | l | x | x | l | l |
| x | l | x | l | l | l | x | l | x | l |
| x | l | x | l | l | l | x | x | l | l |
| x | l | x | x | l | l | l | l | x | l |
| x | l | x | x | h | x | h | h | h | h |
| x | x | h | l | l | l | x | l | x | l |
| x | x | l | l | l | l | x | h | x | l |
| x | x | h | l | l | l | x | x | l | l |
| x | x | l | x | l | l | l | l | x | l |
| x | x | x | l | l | l | l | l | x | l |
| x | x | x | l | l | l | x | h | l | l |
| x | x | x | h | l | x | h | h | h | h |
| x | l | l | l | l | x | l | x | l | l |
| x | x | x | x | x | l | l | h | l | x |
| x | x | x | x | x | l | l | x | h | h |
| l | x | x | x | x | l | l | x | x | h |
| l | x | x | x | x | l | x | h | l | x |
| l | x | x | x | x | l | x | l | l | x |
| l | x | x | x | x | l | x | x | l | h |
| l | x | x | x | x | x | l | h | x | l |
| l | x | x | x | x | x | l | l | x | l |
| l | h | x | x | x | l | x | x | l | x |
| l | l | x | x | x | l | x | x | l | x |
| l | l | x | x | x | l | x | x | x | h |
| l | l | x | x | x | x | l | x | x | l |
| l | l | x | x | x | x | x | x | l | l |
| l | h | l | x | x | l | x | x | x | x |
| l | l | h | x | x | x | h | x | x | x |
| l | l | h | x | l | x | x | x | x | x |
| l | l | l | x | h | x | x | x | x | x |
| l | l | x | h | x | x | l | x | x | x |
| l | l | x | h | x | x | x | h | x | x |
| l | l | x | h | h | x | x | x | x | x |

| | | | | | | | | | |
|---|---|---|---|---|---|---|---|---|---|
| l | h | x | x | l | x | x | x | l | x |
| l | l | x | x | h | x | x | x | h | x |
| l | x | h | x | x | l | x | l | x | x |
| l | x | l | x | x | l | x | x | l | x |
| l | x | l | x | x | l | x | x | x | h |
| l | x | l | x | x | l | x | x | x | l |
| l | x | h | x | x | x | h | l | x | x |
| l | x | l | x | x | x | l | x | h | x |
| l | x | l | h | x | l | x | x | x | x |
| l | x | h | h | x | x | x | l | x | x |
| l | x | h | l | x | x | x | l | x | x |
| l | x | h | l | l | x | x | x | x | x |
| l | x | h | x | l | x | x | x | h | x |
| l | x | x | l | x | l | x | l | x | x |
| l | x | x | h | x | x | l | l | x | x |
| l | x | x | h | x | x | x | h | x | l |
| l | x | x | l | x | x | x | l | x | l |
| l | x | x | h | l | l | x | x | x | x |
| l | x | x | l | h | x | x | x | x | l |
| l | x | x | x | h | l | l | x | x | x |
| l | x | x | x | l | l | x | x | x | l |
| l | x | x | x | h | x | l | x | h | x |
| l | x | x | x | l | x | l | x | x | h |
| l | x | x | x | h | x | x | l | x | l |
| l | x | x | x | l | x | x | h | x | l |
| x | l | l | x | x | x | l | x | h | x |
| x | l | l | h | h | x | x | x | x | x |
| x | l | x | h | x | l | x | x | l | x |
| x | l | x | h | x | x | l | x | x | h |
| x | l | x | h | h | x | l | x | x | x |
| x | x | l | x | x | l | x | h | l | x |
| x | x | l | x | x | x | l | x | h | l |
| x | x | l | h | x | l | l | x | x | x |
| x | x | l | h | x | l | x | l | x | x |
| x | x | l | h | x | x | l | l | x | x |
| x | x | l | l | x | x | l | x | h | x |
| x | x | l | x | l | x | l | x | h | x |
| x | x | x | h | x | l | l | x | l | x |
| x | x | x | l | x | l | l | x | h | x |
| x | x | x | h | x | l | x | h | l | x |
| l | x | x | x | x | h | h | x | l | l |
| l | x | x | x | x | h | x | h | l | l |
| l | h | x | x | x | l | h | x | h | x |
| l | h | x | x | x | l | h | x | x | h |
| l | l | x | x | x | h | x | h | x | l |
| l | l | x | x | x | l | x | h | x | l |
| l | l | x | x | x | l | x | l | x | l |
| l | h | x | x | x | l | x | x | h | l |
| l | l | x | x | x | l | x | x | h | l |
| l | l | x | x | x | x | h | h | h | x |
| l | l | x | x | x | x | h | h | x | h |
| l | l | x | x | x | x | h | x | h | h |
| l | l | x | x | x | x | l | x | h | h |

| | | | | | | | | | |
|---|---|---|---|---|---|---|---|---|---|
| l | l | x | x | x | x | x | h | h | h |
| l | l | x | x | x | x | x | l | h | l |
| l | l | l | x | x | x | h | h | x | x |
| l | l | l | x | x | x | l | h | x | x |
| l | h | h | x | x | x | h | x | x | l |
| l | l | l | x | x | x | l | x | x | h |
| l | h | h | x | x | x | x | h | x | l |
| l | h | h | x | x | x | x | x | h | l |
| l | h | h | l | x | h | x | x | x | x |
| l | l | l | l | x | l | x | x | x | x |
| l | h | h | l | x | x | x | h | x | x |
| l | l | l | h | x | x | x | x | h | x |
| l | h | h | h | x | x | x | x | x | l |
| l | h | l | h | x | x | x | x | x | h |
| l | l | l | h | x | x | x | x | x | h |
| l | h | l | h | h | x | x | x | x | x |
| l | h | l | h | l | x | x | x | x | x |
| l | h | h | x | l | x | x | h | x | x |
| l | l | l | x | l | x | x | h | x | x |
| l | l | h | x | h | x | x | x | x | h |
| l | h | x | h | x | l | x | l | x | x |
| l | l | x | h | x | l | x | l | x | x |
| l | l | x | l | x | l | x | h | x | x |
| l | l | x | l | x | l | x | x | h | x |
| l | h | x | h | x | l | x | x | x | l |
| l | l | x | l | x | h | x | x | x | l |
| l | l | x | h | x | x | h | x | h | x |
| l | l | x | l | x | x | x | h | x | l |
| l | h | x | h | x | x | x | x | h | l |
| l | h | x | h | x | x | x | x | l | l |
| l | l | x | h | x | x | x | x | h | l |
| l | l | x | l | x | x | x | x | h | l |
| l | h | x | h | l | x | h | x | x | x |
| l | h | x | h | l | x | x | x | h | x |
| l | l | x | h | l | x | x | x | l | x |
| l | l | x | l | l | x | x | x | x | l |
| l | h | x | x | l | x | h | h | x | x |
| l | l | x | x | h | x | h | h | x | x |
| l | l | x | x | l | x | l | h | x | x |
| l | x | h | x | x | h | h | x | h | x |
| l | x | h | x | x | x | h | x | h | h |
| l | x | l | x | x | x | x | l | l | l |
| l | x | h | l | x | l | x | x | x | l |
| l | x | l | l | x | x | l | l | x | x |
| l | x | l | h | x | x | l | x | x | l |
| l | x | l | h | x | x | x | h | x | h |
| l | x | l | l | x | x | x | h | x | l |
| l | x | h | h | x | x | x | x | h | l |
| l | x | h | l | x | x | x | x | l | l |
| l | x | l | h | x | x | x | x | h | h |
| l | x | l | h | x | x | x | x | h | l |
| l | x | l | h | x | x | x | x | l | l |
| l | x | l | l | x | x | x | x | l | l |

| | | | | | | | | | |
|---|---|---|---|---|---|---|---|---|---|
| l | x | l | l | l | l | x | x | x | x |
| l | x | h | h | l | x | h | x | x | x |
| l | x | l | h | h | x | l | x | x | x |
| l | x | l | h | l | x | l | x | x | x |
| l | x | l | l | l | x | l | x | x | x |
| l | x | l | h | h | x | x | x | h | x |
| l | x | h | h | l | x | x | x | x | h |
| l | x | l | h | h | x | x | x | x | h |
| l | x | l | x | l | x | l | h | x | x |
| l | x | l | x | l | x | l | l | x | x |
| l | x | l | x | h | x | h | x | h | x |
| l | x | l | x | h | x | h | x | x | h |
| l | x | l | x | h | x | x | l | h | x |
| l | x | l | x | h | x | x | x | h | h |
| l | x | x | h | x | l | l | x | h | x |
| l | x | x | h | x | l | h | x | x | h |
| l | x | x | h | x | h | x | h | h | x |
| l | x | x | l | x | l | x | h | h | x |
| l | x | x | h | x | l | x | l | x | l |
| l | x | x | l | x | l | x | h | x | l |
| l | x | x | l | x | l | x | x | l | l |
| l | x | x | h | x | x | l | x | h | h |
| l | x | x | h | l | h | x | h | x | x |
| l | x | x | h | l | h | x | x | h | x |
| l | x | x | l | l | l | x | x | h | x |
| l | x | x | h | l | x | h | h | x | x |
| l | x | x | l | l | x | l | h | x | x |
| l | x | x | h | h | x | x | l | h | x |
| l | x | x | h | l | x | x | h | h | x |
| l | x | x | h | l | x | x | h | l | x |
| l | x | x | x | l | x | l | h | l | x |
| l | x | x | x | h | x | h | x | h | h |
| l | x | x | x | l | x | h | x | l | l |
| l | x | x | x | l | x | x | l | l | l |
| x | l | x | x | x | l | x | l | h | l |
| x | l | h | x | x | l | l | x | x | h |
| x | l | l | x | x | l | x | l | x | l |
| x | l | l | x | x | l | x | x | l | l |
| x | l | h | x | x | x | h | x | h | h |
| x | l | h | h | x | x | h | x | x | h |
| x | l | l | h | x | x | x | h | x | h |
| x | l | h | h | x | x | x | x | h | h |
| x | l | l | h | x | x | x | x | h | h |
| x | h | l | x | l | l | x | h | x | x |
| x | h | l | x | l | l | x | x | h | x |
| x | l | l | x | h | l | x | x | h | x |
| x | l | l | x | h | l | x | x | x | h |
| x | l | x | h | x | l | x | l | h | x |
| x | l | x | h | x | l | x | l | x | l |
| x | l | x | l | x | x | l | l | h | x |
| x | l | x | h | x | x | h | h | x | h |
| x | l | x | h | h | l | x | l | x | x |
| x | l | x | h | h | x | x | h | x | h |

| | | | | | | | | | |
|---|---|---|---|---|---|---|---|---|---|
| x | h | x | x | l | l | h | x | h | x |
| x | l | x | x | l | l | l | x | h | x |
| x | l | x | x | h | l | l | x | x | l |
| x | l | x | x | h | l | x | l | h | x |
| x | l | x | x | h | l | x | x | h | h |
| x | l | x | x | l | l | x | x | l | l |
| x | x | l | x | x | l | l | l | x | l |
| x | x | l | l | x | l | x | l | h | x |
| x | x | h | l | l | l | x | l | x | x |
| x | x | l | h | l | l | x | h | x | x |
| x | x | l | l | l | l | x | h | x | x |
| x | x | h | l | l | l | x | x | x | l |
| x | x | l | h | h | x | l | x | h | x |
| x | x | l | l | l | x | x | l | x | l |
| x | x | l | h | l | x | x | x | h | h |
| x | x | h | x | l | l | l | x | l | x |
| x | x | l | x | l | l | l | x | l | x |
| x | x | l | x | h | l | l | x | x | l |
| x | x | l | x | l | l | l | x | x | l |
| x | x | l | x | l | l | x | h | x | l |
| x | x | l | x | h | x | x | l | h | l |
| x | x | x | l | x | l | x | l | h | l |
| x | x | x | l | x | x | l | l | h | l |
| x | x | x | l | l | l | x | h | x | l |
| x | x | x | l | l | x | l | l | h | x |
| x | x | x | l | l | x | l | l | x | l |
| x | x | x | h | l | x | h | x | h | h |
| x | x | x | h | l | x | x | h | h | h |
| x | x | x | x | l | l | l | l | x | l |
| x | x | x | x | l | l | x | h | l | l |
| l | h | l | x | x | x | h | h | x | h |
| l | h | l | x | x | x | x | h | h | h |
| l | h | l | h | x | x | x | h | h | x |
| l | l | l | x | l | x | l | x | l | x |
| l | h | x | h | x | l | x | x | h | h |
| l | x | h | h | x | l | l | h | x | x |
| l | x | l | l | l | x | x | l | l | x |
| l | x | x | h | x | x | h | h | h | h |
| l | x | x | l | h | x | h | l | x | h |
| l | x | x | h | l | x | x | l | h | l |
| l | x | x | x | h | h | h | h | h | x |
| x | l | l | x | x | x | l | h | l | l |
| x | l | l | l | x | x | l | h | x | l |
| x | l | l | l | x | x | l | l | x | l |
| x | l | l | l | x | x | x | l | h | l |
| x | l | l | l | l | l | x | x | x | l |
| x | l | l | l | l | x | l | h | x | x |
| x | l | l | l | l | x | x | h | l | x |
| x | l | l | l | l | x | x | x | l | l |
| x | l | h | x | l | l | l | l | x | x |
| x | l | l | x | l | l | l | h | x | x |
| x | l | l | x | l | l | l | l | x | x |
| x | l | l | x | l | x | l | h | l | x |

| | | | | | | | | | |
|---|---|---|---|---|---|---|---|---|---|
| x | l | l | x | l | x | l | l | x | l |
| x | l | l | x | h | x | h | x | h | h |
| x | l | l | x | l | x | x | h | l | h |
| x | l | l | x | l | x | x | l | l | l |
| x | l | x | l | x | l | x | h | l | l |
| x | h | x | h | l | l | h | h | x | x |
| x | l | x | h | l | l | l | l | x | x |
| x | l | x | l | l | l | l | h | x | x |
| x | l | x | h | h | x | h | h | h | x |
| x | l | x | l | l | x | x | l | l | l |
| x | l | x | x | h | x | l | h | l | l |
| x | l | x | x | l | x | l | h | l | h |
| x | x | h | l | h | l | l | h | x | x |
| x | x | l | l | l | l | l | l | x | x |
| x | x | l | l | l | l | x | x | h | l |
| x | x | l | l | l | x | l | x | l | l |
| x | x | h | l | l | x | x | l | l | l |
| x | x | l | x | h | l | h | l | x | h |
| x | l | h | x | l | x | l | l | l | l |
| l | x | x | x | x | x | l | x | x | l |
| l | x | h | x | x | x | x | l | x | x |
| x | x | x | x | x | l | l | l | h | x |
| l | x | x | x | x | l | l | x | h | x |
| l | x | x | x | x | h | x | h | x | l |
| l | x | x | x | x | l | x | l | x | l |
| l | x | x | x | x | l | x | x | h | l |
| l | x | x | x | x | l | x | x | l | l |
| l | x | x | x | x | x | l | x | h | h |
| l | x | x | x | x | x | x | h | h | l |
| l | x | x | x | x | x | x | h | l | l |
| l | x | x | x | x | x | x | l | l | l |
| l | l | x | x | x | l | x | l | x | x |
| l | h | x | x | x | l | x | x | x | l |
| l | l | x | x | x | h | x | x | x | l |
| l | l | x | x | x | x | h | h | x | x |
| l | h | x | x | x | x | x | h | l | x |
| l | l | x | x | x | x | x | h | l | x |
| l | l | x | x | x | x | x | h | x | l |
| l | l | x | x | x | x | x | x | h | l |
| l | l | h | x | x | l | x | x | x | x |
| l | l | h | x | x | x | l | x | x | x |
| l | l | h | x | x | x | x | x | h | x |
| l | h | h | x | x | x | x | x | x | l |
| l | l | h | x | x | x | x | x | x | l |
| l | h | h | x | l | x | x | x | x | x |
| l | h | l | x | l | x | x | x | x | x |
| l | l | x | h | x | h | x | x | x | x |
| l | l | x | h | x | x | x | x | h | x |
| l | l | x | h | x | x | x | x | l | x |
| l | h | x | h | x | x | x | x | x | l |
| l | l | x | h | x | x | x | x | x | h |
| l | l | x | l | x | x | x | x | x | l |
| l | h | x | h | l | x | x | x | x | x |

| | | | | | | | | | |
|---|---|---|---|---|---|---|---|---|---|
| l | h | x | l | l | x | x | x | x | x |
| l | l | x | x | h | l | x | x | x | x |
| l | l | x | x | h | x | h | x | x | x |
| l | h | x | x | l | x | x | h | x | x |
| l | h | x | x | l | x | x | l | x | x |
| l | l | x | x | l | x | x | h | x | x |
| l | h | x | x | l | x | x | x | x | l |
| l | x | l | x | x | x | x | h | l | x |
| l | x | l | x | x | x | x | h | x | l |
| l | x | h | x | x | x | x | x | h | l |
| l | x | l | l | x | l | x | x | x | x |
| l | x | h | l | x | x | l | x | x | x |
| l | x | h | h | x | x | x | x | x | l |
| l | x | h | l | x | x | x | x | x | l |
| l | x | h | h | l | x | x | x | x | x |
| l | x | l | h | h | x | x | x | x | x |
| l | x | h | x | l | h | x | x | x | x |
| l | x | h | x | l | x | h | x | x | x |
| l | x | l | x | h | x | h | x | x | x |
| l | x | l | x | h | x | x | l | x | x |
| l | x | l | x | h | x | x | x | h | x |
| l | x | x | h | x | l | l | x | x | x |
| l | x | x | h | x | l | x | l | x | x |
| l | x | x | l | x | l | x | h | x | x |
| l | x | x | l | x | l | x | x | h | x |
| l | x | x | h | x | l | x | x | x | l |
| l | x | x | l | x | l | x | x | x | h |
| l | x | x | l | x | l | x | x | x | l |
| l | x | x | h | x | x | l | x | h | x |
| l | x | x | h | x | x | x | l | h | x |
| l | x | x | h | x | x | x | x | h | l |
| l | x | x | h | x | x | x | x | l | l |
| l | x | x | l | x | x | x | x | h | l |
| l | x | x | l | h | l | x | x | x | x |
| l | x | x | l | l | l | x | x | x | x |
| l | x | x | h | l | x | l | x | x | x |
| l | x | x | l | l | x | l | x | x | x |
| l | x | x | h | h | x | x | l | x | x |
| l | x | x | h | l | x | x | h | x | x |
| l | x | x | h | l | x | x | x | x | h |
| l | x | x | l | l | x | x | x | x | l |
| l | x | x | x | l | l | x | l | x | x |
| l | x | x | x | l | l | x | x | h | x |
| l | x | x | x | l | x | l | h | x | x |
| l | x | x | x | l | x | l | l | x | x |
| l | x | x | x | l | x | l | x | h | x |
| l | x | x | x | l | x | x | h | l | x |
| l | x | x | x | h | x | x | h | x | l |
| l | x | x | x | h | x | x | x | l | l |
| l | x | x | x | l | x | x | x | l | l |
| x | l | x | x | x | l | l | x | x | h |
| x | l | x | x | x | x | l | x | h | h |
| x | l | h | x | x | l | x | x | x | h |

| | | | | | | | | | |
|---|---|---|---|---|---|---|---|---|---|
| x | l | l | h | x | x | x | h | x | x |
| x | l | l | h | x | x | x | x | l | x |
| x | h | x | l | x | l | l | x | x | x |
| x | l | x | h | x | l | x | l | x | x |
| x | l | x | h | x | x | x | h | x | h |
| x | l | x | h | h | l | x | x | x | x |
| x | l | x | x | h | l | l | x | x | x |
| x | l | x | x | h | x | l | x | x | l |
| x | x | l | x | x | l | l | x | x | h |
| x | x | l | x | x | l | x | l | h | x |
| x | x | l | l | x | l | x | h | x | x |
| x | x | l | x | l | l | l | x | x | x |
| x | x | l | x | l | l | x | h | x | x |
| x | x | l | x | h | x | l | x | h | x |
| x | x | l | x | l | x | l | x | x | h |
| x | x | x | l | x | l | l | x | x | h |
| x | x | x | l | x | l | x | l | h | x |
| x | x | x | h | x | l | x | x | l | h |
| x | x | x | l | x | x | l | l | h | x |
| x | x | x | h | x | x | l | x | h | h |
| x | x | x | l | x | x | x | l | h | l |
| x | x | x | l | l | l | l | x | x | x |
| x | x | x | l | l | l | x | x | h | x |
| x | x | x | h | l | x | l | x | x | h |
| x | x | x | h | l | x | x | x | h | h |
| x | x | x | x | h | l | l | x | h | x |
| x | x | x | x | l | l | l | x | l | x |
| x | x | x | x | h | l | l | x | x | l |
| x | x | x | x | l | l | l | x | x | h |
| x | x | x | x | l | x | l | x | h | h |
| l | x | x | x | x | l | h | x | h | h |
| l | l | x | x | x | h | h | x | h | x |
| l | l | x | x | x | h | x | l | h | x |
| l | l | x | x | x | l | x | h | h | x |
| l | h | x | x | x | l | x | h | x | h |
| l | h | x | x | x | l | x | x | h | h |
| l | h | x | x | x | x | h | h | x | l |
| l | l | l | x | x | x | l | l | x | x |
| l | h | h | x | x | x | h | x | h | x |
| l | l | l | x | x | x | l | x | l | x |
| l | l | h | x | x | x | x | h | x | h |
| l | l | l | x | x | x | x | h | x | h |
| l | h | l | x | x | x | x | x | h | h |
| l | h | h | l | x | x | h | x | x | x |
| l | l | l | l | x | x | x | h | x | x |
| l | h | l | h | x | x | x | x | h | x |
| l | l | l | h | x | x | x | x | x | l |
| l | l | l | x | l | x | l | x | x | x |
| l | h | l | x | h | x | x | x | x | h |
| l | h | x | h | x | l | h | x | x | x |
| l | h | x | h | x | l | x | x | x | h |
| l | l | x | l | x | x | l | l | x | x |
| l | l | x | l | x | x | x | l | h | x |

| | | | | | | | | | |
|---|---|---|---|---|---|---|---|---|---|
| l | l | x | h | x | x | x | l | x | l |
| l | l | x | l | h | x | x | x | l | x |
| l | l | x | h | l | x | x | x | x | l |
| l | h | x | x | h | h | h | x | x | x |
| l | h | x | x | h | x | h | x | h | x |
| l | l | x | x | h | x | l | x | l | x |
| l | l | x | x | l | x | l | x | l | x |
| l | l | x | x | h | x | x | h | x | h |
| l | x | h | x | x | l | h | x | h | x |
| l | x | h | x | x | h | h | x | x | l |
| l | x | h | x | x | x | h | h | h | x |
| l | x | h | x | x | x | l | h | l | x |
| l | x | h | x | x | x | h | h | x | l |
| l | x | l | x | x | x | l | l | x | h |
| l | x | l | x | x | x | x | l | h | l |
| l | x | h | h | x | h | x | x | h | x |
| l | x | h | l | x | l | x | x | l | x |
| l | x | l | l | x | x | l | h | x | x |
| l | x | l | h | x | x | l | x | l | x |
| l | x | l | h | x | x | l | x | x | h |
| l | x | l | h | x | x | x | h | h | x |
| l | x | h | l | x | x | x | x | h | h |
| l | x | l | h | l | x | x | x | h | x |
| l | x | l | h | l | x | x | x | l | x |
| l | x | x | h | x | h | h | x | h | x |
| l | x | x | h | x | h | h | x | x | l |
| l | x | x | h | x | h | x | x | h | h |
| l | x | x | h | x | l | x | x | h | h |
| l | x | x | h | x | x | h | h | h | x |
| l | x | x | l | x | x | l | h | l | x |
| l | x | x | h | x | x | h | x | h | h |
| l | x | x | h | l | h | h | x | x | x |
| l | x | x | l | h | x | h | l | x | x |
| l | x | x | l | l | x | h | x | l | x |
| l | x | x | l | h | x | h | x | x | h |
| l | x | x | l | l | x | x | l | l | x |
| l | x | x | x | h | h | h | x | h | x |
| l | x | x | x | h | l | h | x | h | x |
| l | x | x | x | h | l | h | x | x | h |
| l | x | x | x | h | l | x | x | h | h |
| l | x | x | x | h | x | h | h | h | x |
| l | x | x | x | h | x | h | l | h | x |
| l | x | x | x | h | x | h | l | x | h |
| l | x | x | x | l | x | x | l | h | l |
| x | l | x | x | x | l | h | l | l | x |
| x | l | x | x | x | l | l | l | x | l |
| x | l | x | x | x | l | h | x | h | h |
| x | l | x | x | x | l | l | x | h | l |
| x | l | x | x | x | l | l | x | l | l |
| x | l | x | x | x | l | x | h | l | l |
| x | l | x | x | x | x | l | h | l | l |
| x | l | h | x | x | l | h | x | h | x |
| x | l | l | x | x | l | h | x | l | x |

| | | | | | | | | | |
|---|---|---|---|---|---|---|---|---|---|
| x | l | l | x | x | l | l | x | x | l |
| x | h | l | x | x | l | x | x | h | l |
| x | l | l | x | x | x | l | h | x | l |
| x | l | l | x | x | x | x | l | h | l |
| x | l | l | h | x | l | x | x | x | l |
| x | l | l | h | x | x | x | l | x | l |
| x | l | l | l | x | x | x | l | x | l |
| x | h | h | l | l | l | x | x | x | x |
| x | l | l | h | l | l | x | x | x | x |
| x | l | h | h | l | x | h | x | x | x |
| x | l | h | h | h | x | x | x | x | h |
| x | l | l | x | l | l | x | l | x | x |
| x | l | h | x | h | l | x | x | h | x |
| x | l | h | x | l | l | x | x | l | x |
| x | l | l | x | h | l | x | x | x | l |
| x | l | l | x | l | x | l | h | x | x |
| x | l | l | x | h | x | x | l | h | x |
| x | l | l | x | l | x | x | h | l | x |
| x | l | h | x | h | x | x | x | h | h |
| x | l | x | h | x | l | l | x | h | x |
| x | l | x | l | x | l | x | h | x | l |
| x | l | x | l | x | l | x | l | x | l |
| x | l | x | l | l | l | x | h | x | x |
| x | l | x | l | l | l | x | x | x | l |
| x | l | x | h | h | x | h | x | h | x |
| x | l | x | h | h | x | h | x | x | h |
| x | l | x | l | l | x | x | l | x | l |
| x | l | x | h | h | x | x | x | h | h |
| x | l | x | l | l | x | x | x | l | l |
| x | l | x | x | l | l | x | l | x | l |
| x | l | x | x | l | x | l | h | l | x |
| x | l | x | x | h | x | h | x | h | h |
| x | l | x | x | h | x | x | h | h | h |
| x | x | h | x | x | l | l | l | x | l |
| x | x | l | x | x | l | l | h | x | l |
| x | x | h | x | x | l | l | x | l | l |
| x | x | l | x | x | l | l | x | l | l |
| x | x | l | h | x | l | x | h | x | l |
| x | x | l | h | x | l | x | x | h | l |
| x | x | l | l | x | x | l | l | x | l |
| x | x | h | h | l | l | h | x | x | x |
| x | x | h | l | h | l | l | x | x | x |
| x | x | h | l | l | l | x | x | l | x |
| x | x | l | h | l | l | x | x | h | x |
| x | x | l | h | l | x | x | h | h | x |
| x | x | h | l | l | x | x | l | x | l |
| x | x | l | h | h | x | x | x | h | l |
| x | x | l | l | l | x | x | x | l | l |
| x | x | l | x | l | l | x | x | h | l |
| x | x | l | x | l | x | l | l | x | l |
| x | x | x | l | x | l | l | h | x | l |
| x | x | x | l | x | l | x | h | l | l |
| x | x | x | l | l | l | x | l | x | l |

| | | | | | | | | | |
|---|---|---|---|---|---|---|---|---|---|
| x | x | x | l | l | l | x | x | l | l |
| x | x | x | l | l | x | l | h | x | h |
| x | x | x | l | l | x | x | l | l | l |
| x | x | x | x | l | l | h | h | x | l |
| x | x | x | x | h | l | x | h | l | h |
| x | x | x | x | h | x | l | h | l | l |
| x | x | x | x | l | x | l | h | l | h |
| l | l | l | l | x | x | x | l | l | x |
| l | l | l | l | l | x | x | l | x | x |
| l | l | l | l | l | x | x | x | l | x |
| l | x | l | h | x | h | h | h | x | x |
| x | l | l | x | x | l | l | l | l | x |
| x | h | l | x | x | l | h | l | x | h |
| x | l | h | l | x | l | l | h | x | x |
| x | l | l | l | x | l | l | l | x | x |
| x | l | l | l | x | x | l | x | l | l |
| x | l | l | l | l | x | l | x | x | l |
| x | l | l | x | h | x | h | h | x | h |
| x | l | h | x | l | x | l | x | l | l |
| x | l | h | x | l | x | x | l | l | l |
| x | h | x | h | l | x | h | h | h | x |
| x | l | x | x | l | x | l | l | h | l |
| x | x | x | h | l | l | h | h | h | x |
| x | x | x | x | x | x | l | x | h | h |
| l | x | x | x | x | l | l | x | x | x |
| l | x | x | x | x | x | x | x | h | l |
| l | h | x | x | l | x | x | x | x | x |
| l | x | h | x | l | x | x | x | x | x |
| l | x | x | h | x | x | x | x | x | l |
| l | x | x | x | l | l | x | x | x | x |
| x | x | l | x | x | x | l | x | h | x |
| x | x | x | x | l | l | x | x | x | h |
| x | x | x | x | l | x | l | x | x | h |
| x | x | x | x | x | l | l | h | x | h |
| x | x | x | x | x | l | l | l | x | l |
| x | x | x | x | x | l | l | x | l | h |
| x | x | x | x | x | l | l | x | l | l |
| x | x | x | x | x | l | x | h | l | l |
| l | x | x | x | x | h | h | x | x | l |
| l | x | x | x | x | l | h | x | x | h |
| l | x | x | x | x | l | x | l | h | x |
| l | x | x | x | x | l | x | l | x | h |
| l | x | x | x | x | h | x | x | l | l |
| l | x | x | x | x | l | x | x | h | h |
| l | x | x | x | x | x | l | h | l | x |
| l | h | x | x | x | h | h | x | x | x |
| l | h | x | x | x | l | h | x | x | x |
| l | h | x | x | x | l | x | h | x | x |
| l | l | x | x | x | h | x | h | x | x |
| l | l | x | x | x | l | x | h | x | x |
| l | l | x | x | x | l | x | x | h | x |
| h | l | x | x | x | l | x | x | x | h |
| l | h | x | x | x | l | x | x | x | h |

| | | | | | | | | | |
|---|---|---|---|---|---|---|---|---|---|
| l | l | x | x | x | l | x | x | x | l |
| l | h | x | x | x | x | h | h | x | x |
| l | l | x | x | x | x | l | l | x | x |
| l | l | x | x | x | x | h | x | h | x |
| l | l | x | x | x | x | l | x | h | x |
| l | l | x | x | x | x | l | x | l | x |
| l | h | x | x | x | x | x | l | h | x |
| l | h | x | x | x | x | x | h | x | l |
| l | h | x | x | x | x | x | l | x | l |
| l | l | x | x | x | x | x | h | x | h |
| l | l | x | x | x | x | x | l | x | l |
| l | h | x | x | x | x | x | x | l | l |
| l | l | l | x | x | x | l | x | x | x |
| l | h | l | x | x | x | x | h | x | x |
| l | l | l | x | x | x | x | h | x | x |
| l | h | h | x | x | x | x | x | l | x |
| l | l | h | x | x | x | x | x | l | x |
| l | l | h | x | x | x | x | x | x | h |
| l | l | l | x | x | x | x | x | x | l |
| l | h | l | h | x | x | x | x | x | x |
| l | h | l | x | h | x | x | x | x | x |
| l | l | h | x | h | x | x | x | x | x |
| l | l | x | h | x | l | x | x | x | x |
| l | l | x | l | x | l | x | x | x | x |
| l | l | x | l | x | x | h | x | x | x |
| l | h | x | l | x | x | x | h | x | x |
| l | h | x | l | x | x | x | l | x | x |
| l | h | x | h | x | x | x | x | l | x |
| l | h | x | l | x | x | x | x | x | l |
| l | l | x | h | l | x | x | x | x | x |
| l | l | x | l | h | x | x | x | x | x |
| l | l | x | x | h | x | l | x | x | x |
| l | l | x | x | l | x | l | x | x | x |
| l | l | x | x | h | x | x | x | l | x |
| l | l | x | x | h | x | x | x | x | h |
| l | l | x | x | l | x | x | x | x | l |
| l | x | l | x | x | l | x | l | x | x |
| l | x | h | x | x | h | x | x | h | x |
| l | x | h | x | x | l | x | x | l | x |
| l | x | l | x | x | l | x | x | h | x |
| l | x | l | x | x | x | l | l | x | x |
| l | x | h | x | x | x | h | x | h | x |
| l | x | h | x | x | x | l | x | l | x |
| l | x | h | x | x | x | h | x | x | l |
| l | x | h | x | x | x | x | x | l | l |
| l | x | l | x | x | x | x | x | l | l |
| l | x | h | l | x | l | x | x | x | x |
| l | x | l | h | x | x | l | x | x | x |
| l | x | l | l | x | x | l | x | x | x |
| l | x | h | l | x | x | x | x | h | x |
| l | x | l | l | x | x | x | x | x | l |
| l | x | l | x | l | x | x | h | x | x |
| l | x | x | h | x | h | x | x | h | x |

| | | | | | | | | | |
|---|---|---|---|---|---|---|---|---|---|
| l | x | x | l | x | l | x | x | l | x |
| l | x | x | h | x | l | x | x | x | h |
| l | x | x | l | x | h | x | x | x | l |
| l | x | x | h | x | x | l | x | l | x |
| l | x | x | l | x | x | l | x | h | x |
| l | x | x | h | x | x | x | h | l | x |
| l | x | x | h | x | x | x | l | l | x |
| l | x | x | l | x | x | x | h | x | l |
| l | x | x | l | x | x | x | x | l | l |
| l | x | x | h | l | h | x | x | x | x |
| l | x | x | l | l | x | x | h | x | x |
| l | x | x | h | l | x | x | x | h | x |
| l | x | x | h | l | x | x | x | l | x |
| l | x | x | x | h | h | h | x | x | x |
| l | x | x | x | l | h | x | h | x | x |
| l | x | x | x | h | l | x | x | h | x |
| l | x | x | x | h | l | x | x | l | x |
| l | x | x | x | h | l | x | x | x | l |
| l | x | x | x | h | x | h | l | x | x |
| l | x | x | x | h | x | h | x | h | x |
| l | x | x | x | l | x | l | x | l | x |
| l | x | x | x | l | x | h | x | x | h |
| l | x | x | x | l | x | h | x | x | l |
| l | x | x | x | h | x | x | l | h | x |
| l | x | x | x | h | x | x | l | l | x |
| l | x | x | x | l | x | x | h | x | h |
| x | h | x | x | x | l | l | x | l | x |
| x | l | x | x | x | l | l | x | h | x |
| x | l | x | x | x | l | h | x | x | h |
| x | l | x | x | x | l | x | l | h | x |
| x | l | x | x | x | l | x | x | h | h |
| x | l | x | x | x | l | x | x | l | h |
| x | l | l | x | x | l | x | l | x | x |
| x | l | l | x | x | x | x | l | x | l |
| x | l | l | h | x | l | x | x | x | x |
| x | l | h | h | x | x | x | x | x | h |
| x | l | l | h | x | x | x | x | x | h |
| x | l | l | x | h | l | x | x | x | x |
| x | l | l | x | h | x | x | x | x | l |
| x | l | x | h | x | l | l | x | x | x |
| x | l | x | l | x | l | x | h | x | x |
| x | l | x | l | x | l | x | x | h | x |
| x | l | x | h | x | x | h | x | x | h |
| x | l | x | h | x | x | x | x | h | h |
| x | h | x | l | l | l | x | x | x | x |
| x | l | x | h | h | x | x | h | x | x |
| x | l | x | h | h | x | x | x | x | h |
| x | h | x | x | l | l | h | x | x | x |
| x | h | x | x | l | l | l | x | x | x |
| x | l | x | x | l | l | l | x | x | x |
| x | h | x | x | l | l | x | h | x | x |
| x | l | x | x | l | l | x | l | x | x |
| x | h | x | x | l | l | x | x | h | x |

| | | | | | | | | | |
|---|---|---|---|---|---|---|---|---|---|
| x | l | x | x | h | l | x | x | x | h |
| x | x | h | x | x | l | l | x | x | h |
| x | x | l | x | x | l | h | x | x | h |
| x | x | l | x | x | l | l | x | x | l |
| x | x | l | x | x | l | x | h | x | l |
| x | x | l | x | x | l | x | x | h | l |
| x | x | l | x | x | x | x | l | h | l |
| x | x | h | l | x | l | x | x | h | x |
| x | x | l | l | x | l | x | x | h | x |
| x | x | l | h | x | l | x | x | x | l |
| x | x | h | l | l | l | x | x | x | x |
| x | x | l | h | l | l | x | x | x | x |
| x | x | l | h | h | x | l | x | x | x |
| x | x | l | l | l | x | l | x | x | x |
| x | x | l | x | h | l | l | x | x | x |
| x | x | l | x | l | l | x | l | x | x |
| x | x | l | x | h | x | l | x | x | l |
| x | x | l | x | h | x | x | x | h | l |
| x | x | x | h | x | l | l | x | x | h |
| x | x | x | h | x | l | x | l | l | x |
| x | x | x | h | h | l | l | x | x | x |
| x | x | x | h | l | l | l | x | x | x |
| x | x | x | l | h | l | l | x | x | x |
| x | x | x | h | h | l | x | l | x | x |
| x | x | x | l | l | l | x | h | x | x |
| x | x | x | h | l | l | x | x | l | x |
| x | x | x | l | l | l | x | x | x | l |
| x | x | x | h | h | x | l | x | h | x |
| x | x | x | l | l | x | x | l | x | l |
| x | x | x | x | l | l | l | x | h | x |
| x | x | x | x | h | l | l | x | x | h |
| x | x | x | x | h | l | x | h | l | x |
| x | x | x | x | l | l | x | h | l | x |
| x | x | x | x | l | l | x | x | l | l |
| x | x | x | x | h | x | l | h | x | l |
| l | x | x | x | x | h | h | h | l | x |
| l | x | x | x | x | x | h | h | h | h |
| l | h | x | x | x | x | h | x | h | h |
| l | l | x | x | x | x | x | l | h | h |
| l | l | l | x | x | l | h | x | x | x |
| l | h | l | x | x | x | h | x | x | h |
| l | l | l | x | x | x | h | x | x | h |
| l | l | l | h | x | x | h | x | x | x |
| l | l | l | l | x | x | x | l | x | x |
| l | l | l | l | x | x | x | x | l | x |
| l | h | x | h | x | l | x | x | h | x |
| l | h | x | h | x | x | h | x | h | x |
| l | l | x | l | x | x | x | l | l | x |
| l | h | x | h | x | x | x | l | x | h |
| l | h | x | h | h | l | x | x | x | x |
| l | l | x | l | l | x | x | l | x | x |
| l | h | x | x | h | h | x | x | h | x |
| l | h | x | x | h | x | h | x | x | h |

| | | | | | | | | | |
|---|---|---|---|---|---|---|---|---|---|
| l | h | x | x | h | x | x | h | h | x |
| l | x | h | h | x | h | h | x | x | x |
| l | x | h | h | x | l | x | h | x | x |
| l | x | h | h | x | l | x | x | h | x |
| l | x | h | h | x | x | l | h | x | x |
| l | x | l | h | x | x | h | h | x | x |
| l | x | l | h | x | x | h | x | h | x |
| l | x | h | l | x | x | x | h | l | x |
| l | x | h | h | x | x | x | x | h | h |
| l | x | l | l | l | x | x | l | x | x |
| l | x | l | l | l | x | x | x | l | x |
| l | x | l | x | h | x | l | x | l | x |
| l | x | l | x | l | x | x | l | l | x |
| l | x | x | h | x | l | h | x | h | x |
| l | x | x | l | x | x | l | l | l | x |
| l | x | x | l | x | x | h | l | x | h |
| l | x | x | l | x | x | h | x | l | h |
| l | x | x | h | h | l | h | x | x | x |
| l | x | x | l | l | h | x | l | x | x |
| l | x | x | l | h | x | l | x | l | x |
| l | x | x | l | l | x | x | l | h | x |
| l | x | x | x | h | h | x | h | h | x |
| l | x | x | x | h | x | h | h | x | h |
| x | h | x | x | x | l | h | l | x | h |
| x | l | x | x | x | l | l | h | x | l |
| x | l | x | x | x | l | x | l | l | l |
| x | l | x | x | x | x | l | l | h | l |
| x | l | h | x | x | l | l | x | l | x |
| x | l | l | x | x | l | l | x | l | x |
| x | h | l | x | x | l | x | l | x | h |
| x | l | l | x | x | x | l | h | l | x |
| x | l | h | x | x | x | l | x | l | l |
| x | l | l | x | x | x | h | x | l | l |
| x | l | h | x | x | x | x | h | h | h |
| x | l | l | x | x | x | x | h | l | l |
| x | l | h | h | x | l | h | x | x | x |
| x | l | l | l | x | l | l | x | x | x |
| x | l | h | l | x | l | x | l | x | x |
| x | l | l | l | x | l | x | x | x | l |
| x | l | l | l | x | x | l | h | x | x |
| x | l | h | h | x | x | h | x | h | x |
| x | l | l | h | x | x | h | x | h | x |
| x | l | l | l | x | x | l | x | x | l |
| x | l | h | l | x | x | x | l | x | l |
| x | l | l | h | l | x | x | l | x | x |
| x | l | h | h | h | x | x | x | h | x |
| x | l | l | x | l | l | x | x | l | x |
| x | l | l | x | l | l | x | x | x | l |
| x | l | l | x | h | x | h | x | h | x |
| x | l | l | x | h | x | h | x | x | h |
| x | l | l | x | h | x | x | x | h | h |
| x | l | x | l | x | l | x | x | l | l |
| x | l | x | h | x | x | x | l | l | l |

| | | | | | | | | | |
|---|---|---|---|---|---|---|---|---|---|
| x | h | x | l | l | x | l | l | x | x |
| x | h | x | h | l | x | h | x | h | x |
| x | l | x | h | l | x | h | x | h | x |
| x | h | x | l | l | x | x | l | h | x |
| x | l | x | h | l | x | x | l | l | x |
| x | h | x | l | l | x | x | x | h | l |
| x | l | x | x | l | l | x | h | h | x |
| x | l | x | x | h | l | x | l | x | l |
| x | l | x | x | h | x | h | h | x | h |
| x | l | x | x | l | x | h | h | x | l |
| x | l | x | x | h | x | x | l | h | l |
| x | l | x | x | l | x | x | h | l | h |
| x | l | x | x | l | x | x | l | l | l |
| x | x | h | l | x | l | l | x | l | x |
| x | x | h | l | x | l | l | x | x | l |
| x | x | h | l | x | l | x | l | x | l |
| x | x | l | l | x | l | x | l | x | l |
| x | x | h | l | x | x | l | l | x | l |
| x | x | l | l | x | x | l | h | x | l |
| x | x | l | l | x | x | l | x | l | l |
| x | x | l | h | x | x | x | h | h | l |
| x | x | l | h | l | x | h | h | x | x |
| x | x | l | h | h | x | x | l | h | x |
| x | x | l | x | h | l | x | l | x | h |
| x | x | l | x | h | l | x | x | l | h |
| x | x | l | x | l | x | x | h | l | l |
| x | x | x | h | x | l | h | l | x | h |
| x | x | x | h | l | l | x | l | x | l |
| x | x | x | x | l | l | h | h | h | x |
| x | x | x | x | h | l | h | l | x | h |
| x | x | x | x | h | l | x | l | h | l |
| x | x | x | x | l | l | x | l | h | l |
| l | h | x | h | h | x | x | x | h | h |
| l | x | h | h | h | x | x | h | h | x |
| x | h | h | h | l | h | x | x | h | x |
| x | l | h | l | l | x | x | l | l | x |
| x | l | l | l | l | x | x | l | h | x |
| x | l | l | l | l | x | x | h | x | l |
| x | l | l | l | l | x | x | x | h | l |
| x | l | h | x | l | x | l | l | l | x |
| x | l | h | x | l | x | l | l | x | l |
| x | l | l | x | l | x | l | x | l | l |
| x | l | l | x | l | x | x | h | h | l |
| x | l | x | h | x | h | h | h | h | x |
| x | h | x | h | l | h | x | h | h | x |
| x | l | x | h | l | x | l | l | h | x |
| x | x | h | h | l | x | h | h | h | x |
| x | x | h | l | l | x | l | l | l | x |
| x | x | h | x | l | x | h | h | h | h |
| x | x | h | x | l | x | l | l | l | l |
| x | x | x | x | x | l | l | x | x | h |
| l | x | x | x | x | l | x | l | x | x |
| l | x | x | x | x | l | x | x | l | x |

| | | | | | | | | | |
|---|---|---|---|---|---|---|---|---|---|
| l | x | x | x | x | h | x | x | x | l |
| l | x | x | x | x | l | x | x | x | h |
| l | x | x | x | x | x | l | x | h | x |
| l | x | x | x | x | x | x | h | x | l |
| l | l | x | x | x | x | l | x | x | x |
| l | h | x | x | x | x | x | x | x | l |
| l | l | x | x | x | x | x | x | x | l |
| l | l | h | x | x | x | x | x | x | x |
| l | l | x | x | h | x | x | x | x | x |
| l | x | l | x | x | l | x | x | x | x |
| l | x | h | x | x | x | x | x | x | l |
| l | x | x | h | x | x | x | x | l | x |
| l | x | x | l | x | x | x | x | x | l |
| l | x | x | x | l | x | l | x | x | x |
| l | x | x | x | h | x | x | l | x | x |
| l | x | x | x | l | x | x | h | x | x |
| l | x | x | x | h | x | x | x | x | l |
| l | x | x | x | l | x | x | x | x | l |
| x | l | x | x | x | l | l | x | x | x |
| x | l | x | x | x | l | x | x | x | h |
| x | x | x | l | x | l | x | x | h | x |
| x | x | x | x | l | l | l | x | x | x |
| x | x | x | x | h | x | l | x | x | l |
| x | x | x | x | x | l | l | x | h | l |
| x | x | x | x | x | l | x | h | l | h |
| x | x | x | x | x | l | x | l | h | l |
| l | x | x | x | x | h | h | x | h | x |
| l | x | x | x | x | l | h | x | h | x |
| l | x | x | x | x | x | h | h | l | x |
| l | x | x | x | x | x | h | l | h | x |
| l | x | x | x | x | x | l | l | l | x |
| l | x | x | x | x | x | h | x | h | h |
| l | x | x | x | x | x | h | x | l | l |
| l | l | x | x | x | h | h | x | x | x |
| l | l | x | x | x | l | h | x | x | x |
| l | h | x | x | x | l | x | x | h | x |
| l | h | x | x | x | x | h | x | h | x |
| l | l | x | x | x | x | h | x | x | h |
| l | l | x | x | x | x | x | h | h | x |
| l | l | x | x | x | x | x | l | h | x |
| l | h | x | x | x | x | x | x | l | h |
| l | l | x | x | x | x | x | x | h | h |
| l | h | h | x | x | h | x | x | x | x |
| l | h | l | x | x | x | l | x | x | x |
| l | h | l | x | x | x | x | x | h | x |
| l | l | l | h | x | x | x | x | x | x |
| l | l | l | l | x | x | x | x | x | x |
| l | h | x | h | x | h | x | x | x | x |
| l | h | x | h | x | l | x | x | x | x |
| l | h | x | h | x | x | x | l | x | x |
| l | l | x | h | x | x | x | l | x | x |
| l | l | x | l | x | x | x | h | x | x |
| l | l | x | l | x | x | x | l | x | x |

| | | | | | | | | | |
|---|---|---|---|---|---|---|---|---|---|
| l | h | x | h | x | x | x | x | h | x |
| l | h | x | x | h | l | x | x | x | x |
| l | x | h | x | x | h | h | x | x | x |
| l | x | l | x | x | x | l | h | x | x |
| l | x | l | x | x | x | h | x | x | h |
| l | x | l | x | x | x | l | x | x | h |
| l | x | l | x | x | x | x | h | x | h |
| l | x | h | x | x | x | x | x | h | h |
| l | x | h | h | x | h | x | x | x | x |
| l | x | h | h | x | l | x | x | x | x |
| l | x | l | h | x | x | x | h | x | x |
| l | x | l | l | x | x | x | h | x | x |
| l | x | h | h | x | x | x | x | h | x |
| l | x | h | l | x | x | x | x | l | x |
| l | x | l | h | x | x | x | x | h | x |
| l | x | l | h | x | x | x | x | x | h |
| l | x | l | h | l | x | x | x | x | x |
| l | x | l | x | h | x | l | x | x | x |
| l | x | h | x | h | x | x | x | l | x |
| l | x | l | x | h | x | x | x | l | x |
| l | x | l | x | l | x | x | x | x | h |
| l | x | x | h | x | h | x | h | x | x |
| l | x | x | h | x | l | x | h | x | x |
| l | x | x | h | x | x | h | h | x | x |
| l | x | x | l | x | x | h | l | x | x |
| l | x | x | l | x | x | l | h | x | x |
| l | x | x | l | x | x | l | l | x | x |
| l | x | x | h | x | x | h | x | h | x |
| l | x | x | l | x | x | h | x | l | x |
| l | x | x | l | x | x | l | x | l | x |
| l | x | x | h | x | x | l | x | x | h |
| l | x | x | l | x | x | x | l | h | x |
| l | x | x | h | h | l | x | x | x | x |
| l | x | x | h | h | x | l | x | x | x |
| l | x | x | l | h | x | l | x | x | x |
| l | x | x | l | l | x | h | x | x | x |
| l | x | x | h | l | x | x | l | x | x |
| l | x | x | l | l | x | x | l | x | x |
| l | x | x | l | l | x | x | x | l | x |
| l | x | x | x | l | h | h | x | x | x |
| h | x | x | x | l | l | x | x | l | x |
| l | x | x | x | h | h | x | x | h | x |
| l | x | x | x | h | x | l | h | x | x |
| l | x | x | x | l | x | h | x | l | x |
| l | x | x | x | h | x | x | h | h | x |
| l | x | x | x | l | x | x | l | l | x |
| l | x | x | x | h | x | x | x | h | h |
| l | x | x | x | l | x | x | x | h | h |
| x | l | x | x | x | l | h | x | l | x |
| x | l | x | x | x | l | x | h | l | x |
| x | l | x | x | x | l | x | h | x | l |
| x | l | x | x | x | l | x | l | x | l |
| x | l | x | x | x | l | x | x | h | l |

| | | | | | | | | | |
|---|---|---|---|---|---|---|---|---|---|
| x | l | x | x | x | l | x | x | l | l |
| x | h | l | x | x | l | x | h | x | x |
| x | l | h | x | x | l | x | x | h | x |
| x | l | h | x | x | l | x | x | l | x |
| x | l | l | x | x | l | x | x | x | l |
| x | l | l | x | x | x | l | h | x | x |
| x | l | l | x | x | x | l | x | x | h |
| x | l | l | x | x | x | x | l | h | x |
| x | l | h | x | x | x | x | x | h | h |
| x | l | l | l | x | x | l | x | x | x |
| x | l | l | h | x | x | x | l | x | x |
| x | l | l | h | x | x | x | x | h | x |
| x | h | h | x | l | l | x | x | x | x |
| x | h | l | x | l | l | x | x | x | x |
| x | l | l | x | h | x | l | x | x | x |
| x | l | l | x | h | x | x | x | h | x |
| x | l | x | h | x | h | h | x | x | x |
| x | l | x | h | x | h | x | h | x | x |
| x | l | x | h | x | l | x | h | x | x |
| x | l | x | l | x | l | x | x | x | l |
| x | l | x | h | x | x | l | h | x | x |
| x | l | x | h | x | x | x | l | l | x |
| x | l | x | l | h | l | x | x | x | x |
| x | l | x | l | l | l | x | x | x | x |
| x | l | x | h | h | x | h | x | x | x |
| x | l | x | h | l | x | h | x | x | x |
| x | h | x | l | l | x | x | x | x | l |
| x | l | x | h | l | x | x | x | x | h |
| x | h | x | x | l | l | x | x | l | x |
| x | l | x | x | h | l | x | x | h | x |
| x | l | x | x | l | l | x | x | l | x |
| x | h | x | x | l | l | x | x | x | l |
| x | l | x | x | h | l | x | x | x | l |
| x | x | h | x | x | l | l | x | l | x |
| x | x | h | x | x | l | x | h | l | x |
| x | x | l | x | x | l | x | h | x | h |
| x | x | l | x | x | l | x | l | x | l |
| x | x | l | x | x | l | x | x | l | h |
| x | x | l | x | x | x | l | l | x | l |
| x | x | h | l | x | l | l | x | x | x |
| x | x | l | h | x | x | l | x | l | x |
| x | x | l | h | x | x | l | x | x | h |
| x | x | l | l | x | x | l | x | x | l |
| x | x | l | h | x | x | x | l | h | x |
| x | x | l | h | x | x | x | l | x | l |
| x | x | l | l | x | x | x | l | x | l |
| x | x | l | h | x | x | x | x | h | l |
| x | x | l | h | l | x | x | h | x | x |
| x | x | l | h | l | x | x | x | h | x |
| x | x | h | x | l | l | h | x | x | x |
| x | x | l | x | h | l | x | l | x | x |
| x | x | h | x | l | l | x | x | l | x |
| x | x | l | x | l | l | x | x | h | x |

| | | | | | | | | | |
|---|---|---|---|---|---|---|---|---|---|
| x | x | l | x | l | l | x | x | l | x |
| x | x | l | x | l | l | x | x | x | l |
| x | x | l | x | l | x | l | h | x | x |
| x | x | l | x | h | x | x | l | h | x |
| x | x | l | x | l | x | x | h | x | h |
| x | x | x | h | x | l | l | x | h | x |
| x | x | x | h | x | l | l | x | x | l |
| x | x | x | h | x | l | x | l | x | l |
| x | x | x | l | x | l | x | h | x | l |
| x | x | x | h | x | l | x | x | l | l |
| x | x | x | h | x | x | l | l | x | l |
| x | x | x | l | x | x | l | l | x | l |
| x | x | x | l | x | x | l | x | h | l |
| x | x | x | h | l | l | h | x | x | x |
| x | x | x | h | l | l | x | l | x | x |
| x | x | x | l | l | l | x | l | x | x |
| x | x | x | h | l | l | x | x | h | x |
| x | x | x | h | l | x | l | x | l | x |
| x | x | x | l | l | x | l | x | h | x |
| x | x | x | x | h | l | l | x | l | x |
| x | x | x | x | h | l | x | l | h | x |
| x | x | x | x | l | l | x | l | h | x |
| x | x | x | x | l | l | x | l | l | x |
| x | x | x | x | l | l | x | l | x | l |
| l | h | x | x | x | x | h | l | x | h |
| l | h | x | x | x | x | x | h | h | h |
| l | h | h | x | x | x | h | x | x | h |
| l | h | h | x | x | x | x | h | h | x |
| l | h | l | x | x | x | x | l | x | h |
| l | h | h | l | x | x | x | x | x | h |
| l | l | l | x | l | h | x | x | x | x |
| l | h | h | x | h | x | h | x | x | x |
| l | h | h | x | h | x | x | h | x | x |
| l | l | l | x | l | x | x | x | l | x |
| l | l | x | l | l | h | x | x | x | x |
| l | h | x | h | h | x | h | x | x | x |
| l | h | x | h | h | x | x | h | x | x |
| h | l | x | x | l | x | x | x | l | h |
| l | x | h | x | x | l | x | h | h | x |
| l | x | l | l | x | x | x | l | l | x |
| l | x | h | x | h | x | h | x | x | h |
| l | x | x | h | x | x | x | h | h | h |
| l | x | x | l | x | x | x | h | l | h |
| l | x | x | x | l | h | x | l | h | x |
| x | l | x | x | x | h | h | h | h | x |
| x | l | x | x | x | x | h | h | h | h |
| x | l | l | x | x | l | x | h | h | x |
| x | h | h | x | x | l | x | x | l | l |
| x | l | h | x | x | x | h | h | h | x |
| x | l | h | x | x | x | h | h | x | h |
| x | l | h | x | x | x | x | h | l | l |
| x | l | h | x | x | x | x | l | l | l |
| x | l | l | x | x | x | x | h | h | l |

| | | | | | | | | | |
|---|---|---|---|---|---|---|---|---|---|
| x | l | l | x | x | x | x | h | l | h |
| x | h | l | h | x | l | x | x | x | h |
| x | l | l | l | x | x | x | h | l | x |
| x | h | l | h | x | x | x | h | x | h |
| x | l | h | l | x | x | x | x | l | l |
| x | l | l | l | x | x | x | x | h | l |
| x | l | l | l | x | x | x | x | l | l |
| x | h | l | h | l | x | x | x | x | h |
| x | l | h | l | l | x | x | x | x | l |
| x | l | l | l | l | x | x | x | x | l |
| x | l | h | x | h | l | x | h | x | x |
| x | h | l | x | l | x | l | l | x | x |
| x | l | h | x | l | x | h | h | x | x |
| x | l | h | x | l | x | l | l | x | x |
| x | l | h | x | l | x | l | x | l | x |
| x | l | h | x | l | x | x | l | l | x |
| x | l | h | x | h | x | x | h | x | h |
| x | l | l | x | l | x | x | h | x | l |
| x | l | h | x | l | x | x | x | l | l |
| x | l | l | x | l | x | x | x | l | l |
| x | l | x | h | x | l | h | x | h | x |
| x | l | x | h | x | x | h | h | h | x |
| x | l | x | h | x | x | h | l | h | x |
| x | l | x | h | x | x | l | l | h | x |
| x | l | x | l | x | x | h | l | l | x |
| x | l | x | l | x | x | l | h | x | l |
| x | l | x | l | x | x | l | x | l | l |
| x | h | x | h | l | h | x | x | h | x |
| x | l | x | l | l | h | x | x | x | l |
| x | l | x | l | l | x | l | h | x | x |
| x | l | x | h | l | x | l | x | h | x |
| x | l | x | h | h | x | x | l | h | x |
| x | l | x | h | l | x | x | h | h | x |
| x | l | x | l | l | x | x | h | l | x |
| x | l | x | x | l | h | x | h | x | h |
| x | l | x | x | h | x | h | h | h | x |
| x | l | x | x | l | x | l | h | h | x |
| x | l | x | x | l | x | l | l | h | x |
| x | l | x | x | l | x | l | l | x | l |
| x | l | x | x | l | x | h | x | l | l |
| x | l | x | x | l | x | l | x | l | l |
| x | l | x | x | l | x | x | h | h | l |
| x | l | x | x | l | x | x | h | l | l |
| x | x | l | x | x | l | l | l | l | x |
| x | x | h | x | x | l | h | l | x | h |
| x | x | l | x | x | x | l | h | l | l |
| x | x | h | l | x | l | x | l | l | x |
| x | x | h | l | x | l | x | x | l | l |
| x | x | l | l | x | l | x | x | l | l |
| x | x | h | l | x | x | l | x | l | l |
| x | x | h | l | h | l | x | h | x | x |
| x | x | l | l | h | l | x | x | x | h |
| x | x | h | l | l | x | l | l | x | x |

| | | | | | | | | | |
|---|---|---|---|---|---|---|---|---|---|
| x | x | h | h | l | x | h | x | h | x |
| x | x | h | l | l | x | x | l | l | x |
| x | x | l | h | l | x | x | l | l | x |
| x | x | l | l | l | x | x | l | h | x |
| x | x | h | l | l | x | x | x | l | l |
| x | x | h | x | l | x | l | l | l | x |
| x | x | h | x | l | x | h | h | x | l |
| x | x | h | x | l | x | l | l | x | l |
| x | x | h | x | l | x | h | x | h | h |
| x | x | h | x | l | x | h | x | l | l |
| x | x | l | x | l | x | l | x | l | l |
| x | x | h | x | l | x | x | h | h | h |
| x | x | h | x | l | x | x | l | l | l |
| x | x | x | h | x | l | x | l | h | h |
| x | x | x | h | l | h | h | x | h | x |
| x | x | x | h | h | l | x | x | h | l |
| x | x | x | h | l | x | h | h | h | x |
| x | x | x | h | l | x | h | l | h | x |
| x | x | x | l | l | x | l | l | l | x |
| x | x | x | l | l | x | h | x | l | l |
| x | x | x | x | l | h | h | x | l | l |
| x | x | x | x | l | x | l | l | h | l |
| x | x | x | x | l | x | l | l | l | l |
| h | l | h | l | l | x | x | h | x | x |
| x | h | l | x | x | x | h | h | h | h |
| x | h | h | h | l | x | x | h | h | x |
| x | l | l | l | l | x | x | l | l | x |
| x | h | h | x | l | x | h | h | h | x |
| x | x | h | h | l | h | x | h | h | x |
| x | x | x | x | x | l | l | h | x | x |
| x | x | x | x | x | l | l | x | h | x |
| x | x | x | x | x | l | x | h | l | x |
| l | x | x | x | x | l | x | x | h | x |
| l | x | x | x | x | l | x | x | x | l |
| l | x | x | x | x | x | l | l | x | x |
| l | x | x | x | x | x | h | x | h | x |
| l | x | x | x | x | x | h | x | x | l |
| l | x | x | x | x | x | x | h | l | x |
| l | x | x | x | x | x | x | l | x | l |
| l | x | x | x | x | x | x | x | l | l |
| h | l | x | x | x | l | x | x | x | x |
| l | h | x | x | x | l | x | x | x | x |
| l | l | x | x | x | l | x | x | x | x |
| l | h | x | x | x | x | x | l | x | x |
| l | l | x | x | x | x | x | h | x | x |
| l | h | x | x | x | x | x | x | l | x |
| l | l | x | h | x | x | x | x | x | x |
| l | x | h | x | x | x | l | x | x | x |
| l | x | h | x | x | x | x | x | l | x |
| l | x | l | x | x | x | x | x | x | l |
| l | x | h | l | x | x | x | x | x | x |
| l | x | l | x | h | x | x | x | x | x |
| l | x | x | l | x | l | x | x | x | x |

| | | | | | | | | | |
|---|---|---|---|---|---|---|---|---|---|
| l | x | x | h | x | x | l | x | x | x |
| l | x | x | l | x | x | h | x | x | x |
| l | x | x | l | x | x | l | x | x | x |
| l | x | x | h | x | x | x | l | x | x |
| l | x | x | h | l | x | x | x | x | x |
| h | x | x | x | l | l | x | x | x | x |
| l | x | x | x | l | h | x | x | x | x |
| l | x | x | x | h | x | x | x | h | x |
| l | x | x | x | l | x | x | x | x | h |
| x | l | x | x | x | l | x | l | x | x |
| x | l | l | h | x | x | x | x | x | x |
| x | h | x | l | x | l | x | x | x | x |
| x | l | x | h | x | x | x | x | l | x |
| x | l | x | h | x | x | x | x | x | h |
| x | h | x | x | l | l | x | x | x | x |
| x | l | x | x | h | l | x | x | x | x |
| x | h | x | x | l | x | x | l | x | x |
| x | l | x | x | h | x | x | x | x | l |
| x | x | l | x | x | l | l | x | x | x |
| x | x | l | x | x | l | x | l | x | x |
| x | x | l | x | x | l | x | x | x | h |
| x | x | l | x | x | x | l | x | x | h |
| x | x | l | h | x | x | l | x | x | x |
| x | x | h | x | l | l | x | x | x | x |
| x | x | l | x | l | l | x | x | x | x |
| x | x | l | x | h | x | l | x | x | x |
| x | x | x | h | x | l | l | x | x | x |
| x | x | x | l | x | l | l | x | x | x |
| x | x | x | l | x | l | x | h | x | x |
| x | x | x | h | x | l | x | x | l | x |
| x | x | x | l | x | l | x | x | x | h |
| x | x | x | l | x | x | l | x | h | x |
| x | x | x | h | l | l | x | x | x | x |
| x | x | x | l | l | l | x | x | x | x |
| x | x | x | x | h | l | l | x | x | x |
| x | x | x | x | l | l | h | x | x | x |
| x | x | x | x | l | l | x | h | x | x |
| x | x | x | x | l | l | x | l | x | x |
| x | x | x | x | l | l | x | x | h | x |
| x | x | x | x | l | l | x | x | l | x |
| x | x | x | x | l | l | x | x | x | l |
| x | x | x | x | h | x | l | x | h | x |
| x | x | x | x | x | l | h | l | x | h |
| x | x | x | x | x | x | h | h | l | l |
| x | x | x | x | x | x | l | h | l | l |
| x | x | x | x | x | x | l | l | h | l |
| l | x | x | x | x | x | h | l | l | x |
| l | x | x | x | x | x | h | l | x | h |
| l | h | x | x | x | h | x | x | h | x |
| l | l | x | x | x | h | x | x | h | x |
| l | h | x | x | x | x | h | x | x | h |
| l | h | x | x | x | x | x | h | h | x |
| l | h | x | x | x | x | x | x | h | h |

| | | | | | | | | | |
|---|---|---|---|---|---|---|---|---|---|
| l | l | l | x | x | h | x | x | x | x |
| l | h | h | x | x | x | h | x | x | x |
| l | l | l | x | x | x | h | x | x | x |
| l | h | h | x | x | x | x | h | x | x |
| l | h | h | x | x | x | x | x | h | x |
| l | h | l | x | x | x | x | x | x | h |
| l | l | l | x | x | x | x | x | x | h |
| l | l | x | l | x | h | x | x | x | x |
| l | h | x | h | x | x | h | x | x | x |
| l | h | x | h | x | x | x | h | x | x |
| l | h | x | l | x | x | x | x | h | x |
| l | l | x | l | x | x | x | x | l | x |
| l | l | x | l | l | x | x | x | x | x |
| l | h | x | x | h | h | x | x | x | x |
| l | h | x | x | h | x | h | x | x | x |
| l | l | x | x | l | x | h | x | x | x |
| l | h | x | x | h | x | x | h | x | x |
| h | l | x | x | l | x | x | x | l | x |
| l | x | h | x | x | l | h | x | x | x |
| l | x | l | x | x | h | h | x | x | x |
| l | x | h | x | x | l | x | h | x | x |
| l | x | h | x | x | x | h | h | x | x |
| l | x | l | x | x | x | h | h | x | x |
| l | x | l | x | x | x | h | x | l | x |
| l | x | l | x | x | x | l | x | l | x |
| l | x | l | x | x | x | x | l | h | x |
| l | x | h | h | x | x | h | x | x | x |
| l | x | h | h | x | x | x | h | x | x |
| l | x | l | x | l | x | h | x | x | x |
| l | x | l | x | l | x | x | x | l | x |
| l | x | x | h | x | h | h | x | x | x |
| l | x | x | h | x | l | h | x | x | x |
| l | x | x | h | x | x | x | h | h | x |
| l | x | x | l | x | x | x | l | l | x |
| l | x | x | l | x | x | x | h | x | h |
| l | x | x | h | x | x | x | x | h | h |
| l | x | x | h | h | h | x | x | x | x |
| l | x | x | l | h | x | x | x | l | x |
| l | x | x | x | h | h | x | h | x | x |
| l | x | x | x | h | l | x | h | x | x |
| l | x | x | x | h | x | h | h | x | x |
| l | x | x | x | l | x | h | l | x | x |
| l | x | x | x | h | x | l | x | l | x |
| l | x | x | x | h | x | h | x | x | h |
| l | x | x | x | l | x | x | l | h | x |
| x | h | x | x | x | l | l | x | x | l |
| x | l | x | x | x | l | x | h | h | x |
| x | h | x | x | x | l | x | h | x | l |
| x | h | x | x | x | l | x | l | x | h |
| x | h | x | x | x | l | x | x | h | l |
| x | l | x | x | x | x | l | h | h | x |
| x | l | x | x | x | x | l | l | h | x |
| x | l | x | x | x | x | l | h | x | l |

| | | | | | | | | | |
|---|---|---|---|---|---|---|---|---|---|
| x | l | x | x | x | x | l | l | x | h |
| x | l | x | x | x | x | h | x | l | l |
| x | l | x | x | x | x | x | h | l | l |
| x | l | h | x | x | l | h | x | x | x |
| x | l | h | x | x | l | x | h | x | x |
| x | l | l | x | x | l | x | h | x | x |
| x | h | l | x | x | l | x | x | h | x |
| x | h | l | x | x | l | x | x | l | x |
| x | l | l | x | x | h | x | x | h | x |
| x | l | l | x | x | l | x | x | l | x |
| x | l | l | x | x | x | l | l | x | x |
| x | l | h | x | x | x | h | x | h | x |
| x | l | h | x | x | x | h | x | x | h |
| x | l | l | x | x | x | x | h | l | x |
| x | l | h | x | x | x | x | x | l | l |
| x | l | l | x | x | x | x | x | h | h |
| x | l | l | x | x | x | x | x | h | l |
| x | h | l | h | x | l | x | x | x | x |
| x | l | h | h | x | l | x | x | x | x |
| x | l | h | l | x | l | x | x | x | x |
| x | l | l | l | x | l | x | x | x | x |
| x | l | h | h | x | x | h | x | x | x |
| x | l | h | h | x | x | l | x | x | x |
| x | l | l | l | x | x | x | x | h | x |
| x | h | l | l | x | x | x | x | x | l |
| x | h | h | l | l | x | x | x | x | x |
| x | h | l | h | l | x | x | x | x | x |
| x | l | h | h | h | x | x | x | x | x |
| x | l | h | x | l | x | h | x | x | x |
| x | l | h | x | l | x | x | x | l | x |
| x | l | h | x | l | x | x | x | x | h |
| x | l | l | x | h | x | x | x | x | h |
| x | l | x | h | x | l | h | x | x | x |
| x | l | x | h | x | l | x | x | h | x |
| x | l | x | h | x | l | x | x | x | l |
| x | l | x | h | x | x | h | h | x | x |
| x | l | x | h | x | x | h | l | x | x |
| x | l | x | h | x | x | h | x | h | x |
| x | l | x | h | x | x | l | x | h | x |
| x | l | x | h | x | x | x | l | x | l |
| x | l | x | l | x | x | x | l | x | l |
| x | l | x | l | x | x | x | x | h | l |
| x | h | x | l | l | x | l | x | x | x |
| x | l | x | h | h | x | x | l | x | x |
| x | h | x | l | l | x | x | x | h | x |
| x | l | x | h | h | x | x | x | h | x |
| x | l | x | x | l | x | l | h | x | x |
| x | l | x | x | h | x | h | x | h | x |
| x | l | x | x | l | x | l | x | h | x |
| x | l | x | x | h | x | x | l | h | x |
| x | l | x | x | l | x | x | h | l | x |
| x | l | x | x | h | x | x | x | h | h |
| x | l | x | x | l | x | x | x | l | l |

| | | | | | | | | | |
|---|---|---|---|---|---|---|---|---|---|
| x | x | l | x | x | l | h | x | l | x |
| x | x | h | x | x | l | l | x | x | l |
| x | x | l | x | x | l | x | x | l | l |
| x | x | h | x | x | x | l | l | x | l |
| x | x | l | x | x | x | l | h | x | l |
| x | x | h | x | x | x | l | x | l | l |
| x | x | l | x | x | x | x | h | l | l |
| x | x | h | l | x | l | x | l | x | x |
| x | x | h | l | x | l | x | x | l | x |
| x | x | l | l | x | l | x | x | x | l |
| x | x | l | l | x | x | l | h | x | x |
| x | x | l | h | x | x | x | h | x | h |
| x | x | l | h | x | x | x | x | l | l |
| x | x | l | l | x | x | x | x | l | l |
| x | x | l | h | h | l | x | x | x | x |
| x | x | l | l | h | l | x | x | x | x |
| x | x | l | h | l | x | x | l | x | x |
| x | x | l | x | h | l | x | x | x | l |
| x | x | h | x | l | x | l | l | x | x |
| x | x | l | x | l | x | l | l | x | x |
| x | x | l | x | l | x | x | l | h | x |
| x | x | l | x | l | x | x | l | x | l |
| x | x | h | x | l | x | x | x | h | h |
| x | x | x | h | x | l | x | l | x | h |
| x | x | x | l | x | l | x | l | x | l |
| x | x | x | l | x | l | x | x | l | l |
| x | x | x | h | x | x | l | h | h | x |
| x | x | x | l | x | x | l | x | l | l |
| x | x | x | l | h | l | x | x | l | x |
| x | x | x | h | l | x | l | l | x | x |
| x | x | x | l | h | x | l | h | x | x |
| x | x | x | l | l | x | l | l | x | x |
| x | x | x | h | l | x | x | l | l | x |
| x | x | x | l | l | x | x | l | h | x |
| x | x | x | h | l | x | x | l | x | l |
| x | x | x | l | l | x | x | x | l | h |
| x | x | x | l | l | x | x | x | l | l |
| x | x | x | x | h | l | x | x | l | h |
| x | x | x | x | h | x | l | h | l | x |
| x | x | x | x | l | x | h | l | h | x |
| x | x | x | x | l | x | l | l | h | x |
| x | x | x | x | l | x | l | l | x | l |
| x | x | x | x | l | x | h | x | l | l |
| x | x | x | x | l | x | x | l | l | l |
| l | x | x | x | x | h | x | h | h | h |
| l | l | l | x | x | x | x | l | l | x |
| l | h | h | h | x | x | x | x | x | h |
| l | h | x | h | h | x | x | x | x | h |
| l | x | l | l | x | h | x | l | x | x |
| l | x | l | l | x | x | x | l | x | h |
| l | x | l | l | x | x | x | x | l | h |
| x | l | l | x | x | x | l | x | l | l |
| x | l | l | l | x | x | h | l | x | x |

| | | | | | | | | | |
|---|---|---|---|---|---|---|---|---|---|
| x | l | h | l | x | x | l | x | x | l |
| x | l | l | l | x | x | x | l | l | x |
| x | l | l | l | x | x | x | h | x | l |
| x | l | h | h | l | x | x | h | x | x |
| x | l | h | l | l | x | x | h | x | x |
| x | l | h | l | l | x | x | l | x | x |
| x | l | l | l | l | x | x | l | x | x |
| x | h | h | h | l | x | x | x | h | x |
| x | l | l | x | l | h | l | x | x | x |
| x | h | h | x | l | x | h | x | h | x |
| x | h | h | x | l | x | h | x | x | l |
| x | l | l | x | l | x | l | x | x | l |
| x | h | h | x | l | x | x | h | x | l |
| x | h | l | x | l | x | x | h | x | l |
| x | h | h | x | l | x | x | x | h | l |
| x | h | x | h | x | x | x | h | h | l |
| x | l | x | l | l | h | x | x | l | x |
| x | h | x | l | l | x | h | x | l | x |
| x | l | x | l | l | x | l | x | l | x |
| x | h | x | h | l | x | x | h | h | x |
| x | l | x | l | l | x | x | l | l | x |
| x | l | x | h | l | x | x | h | x | l |
| x | l | x | l | l | x | x | h | x | h |
| x | l | x | l | l | x | x | h | x | l |
| x | x | l | h | x | x | h | x | h | h |
| x | x | h | h | h | x | l | h | x | x |
| x | x | h | h | l | x | h | l | x | x |
| x | x | h | h | l | x | h | x | x | l |
| x | x | h | l | l | x | h | x | x | l |
| x | x | l | l | l | x | x | h | l | x |
| x | x | h | x | l | h | x | h | h | x |
| x | x | h | x | l | x | h | h | h | x |
| x | x | x | h | x | x | l | h | l | h |
| x | x | x | h | l | h | x | h | h | x |
| x | x | x | h | l | x | h | h | x | l |
| x | l | l | l | x | x | h | x | l | h |
| x | l | l | l | l | x | h | x | l | x |
